# Disruption of TARP-ARC interaction stabilizes synaptic AMPA receptors and rescues social isolation-induced fear memory deficits

**DOI:** 10.1101/2024.08.04.606518

**Authors:** Jens Edvard Trygstad Gundersen, Xinrong Wu, Jiaxing Song, Aditya Pokharna, Lorena Benedetti, Aamra Mahboob, Håvard Håvardsholm Nesbø, Tadiwos Feyissa Mergiya, William Edward Louch, Huadong Liu, Suya Sun, Jennifer Lippincott-Schwartz, Ingo H. Greger, Nan-jie Xu, Clive R. Bramham, Hongyu Zhang

**Author notes:** These authors contribute equally (Jens Edvard Trygstad Gundersen, Xinrong Wu, Jiaxing Song).

## Abstract

Synaptic trapping of AMPA receptors (AMPARs) is a key mechanism regulating excitatory synaptic transmission and activity-dependent plasticity underlying learning and memory. Destabilization of synaptic AMPARs is increasingly implicated in cognitive dysfunction across neurological and neuropsychiatric disorders, yet strategies to directly modulate this process *in vivo* remain limited, constraining both mechanistic insight and therapeutic development. Here we present a peptide-based strategy to enhance AMPAR synaptic trapping by targeting the interaction between transmembrane AMPA receptor regulatory proteins (TARPs) and activity-regulated cytoskeleton-associated protein (ARC/Arg3.1). We designed a 21-amino-acid TAT-fused peptide (TARP-pep) mimicking the TARP C-terminal motif that binds the ARC N-lobe. TARP-pep disrupted the TARP-ARC interaction and increased the stabilization of AMPARs at synaptic surfaces in cultured hippocampal neurons. *In vivo*, acute intrahippocampal infusion of TARP-pep enhanced perforant path-evoked synaptic transmission in the rat dentate gyrus (DG) and strengthened the interaction between TARPs and postsynaptic density protein 95 (PSD-95), a key mechanism underlying AMPAR anchoring. Consistent with a long-term potentiation (LTP)-like synaptic state, TARP-pep increased basal phosphorylation of Ca^2+^/calmodulin-dependent protein kinase II (CaMKII) and occluded further chemical LTP (cLTP)-induced increases in phosphorylated CaMKII (p-CaMKII). Notably, a 7-day regimen of daily bilateral DG injections of TARP-pep prevented social isolation (SI)-induced impairment of fear memory in mice. Together, these findings identify the TARP-ARC interaction as a druggable regulator of AMPAR diffusional trapping and highlight synaptic AMPAR stabilization as a promising therapeutic strategy for preserving cognitive function under conditions of circuit vulnerability.

## Introduction

As the principal mediators of fast excitatory synaptic transmission, α-amino-3-hydroxy-5-methyl-4-isoxazolepropionic acid receptors (AMPARs) play a central role in synaptic plasticity, learning, and memory formation^1–3^. Critical for these functions, the nanoscale diffusional trapping of AMPARs at synapses enables precise, energy-efficient, real-time tuning of synaptic strength^4–6^. Dysregulation of this process has been implicated in neurological and neuropsychiatric conditions associated with impaired synaptic function, such as Alzheimer’s disease, Huntington’s disease, and stress-related conditions such as depression^**5,7,8**^. Although the structural basis of this trapping, involving scaffold proteins, cytoskeletal networks, adhesion proteins, and the extracellular matrix, is well appreciated, the dynamic regulatory mechanisms by which AMPARs are reversibly and transiently stabilized at synapses remain poorly understood.

Prevailing models of synaptic strengthening propose that neuronal activity either creates new synaptic trapping slots for AMPARs or unmasks pre-existing ones (the slot-centric view), or promotes the exocytic delivery of AMPARs directly into or near dendritic spines (the vesicle-centric view)^9^. In contrast, the mechanisms of activity-dependent synaptic weakening are far less defined. A key candidate is the immediate-early gene product ARC/Arg3.1 (activity-regulated cytoskeleton-associated protein), an activity-dependent regulator of synaptic plasticity that is rapidly induced by neuronal activity and plays a central role in experience-dependent synaptic remodeling^10,11^. Mediation of postsynaptic AMPAR endocytosis has emerged as a canonical function of ARC^12^. Extending beyond this internalization-centric view, we previously proposed that ARC may also regulate synaptic strength by actively disrupting AMPAR anchoring at the level of the trapping mechanism itself^13^.

AMPARs function as macromolecular complexes with auxiliary subunits such as the transmembrane AMPAR regulatory proteins (TARPs). Among TARPs, the prototypical isoforms γ2 (stargazin) and γ8 are highly expressed in the hippocampus and are critical for AMPAR trafficking, synaptic targeting, and stabilization, largely through their interaction with postsynaptic density protein 95 (PSD-95), a major scaffolding protein^14–16^. TARP γ2 and γ8, which share significant sequence homology, also directly interact with ARC via the ARC N-lobe (residues 207–278), which contains a hydrophobic ligand-binding pocket that mediates interactions with synaptic proteins^8^. Based on this interaction, we proposed that ARC may regulate AMPAR synaptic trapping by preventing PSD-95 from binding to the AMPAR-TARP complex, thereby disrupting trapping in an activity-dependent manner^13^ (Fig. 1a, top panel). Consistent with this model, ARC-driven liquid–liquid phase separation can disperse TARPs from PSD condensates^17^. However, direct *in vivo* tests of this ARC–TARP–PSD-95 competition mechanism and its potential therapeutic relevance have been limited by the lack of tools capable of selectively manipulating this specific interaction.

**Fig. 1.**
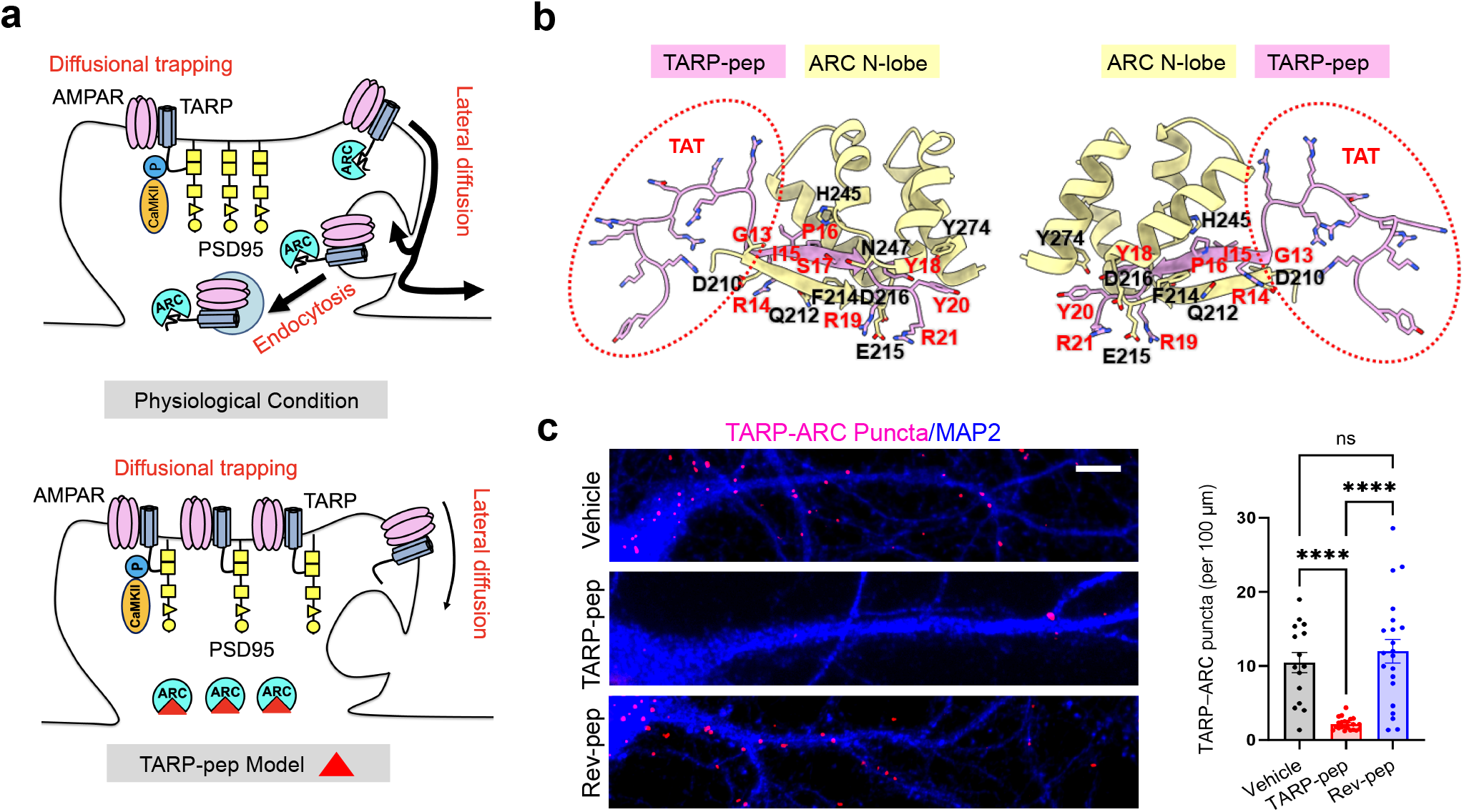
TARP-pep disrupts the TARP-ARC interaction by competitively binding to ARC. **a**, Working model illustrating AMPAR diffusional trapping under physiological conditions (top) and enhanced trapping with TARP-pep (bottom). **b**, Front (left) and back (right) views of the AlphaFold-3-predicted TARP-pep (pink) / ARC N-lobe (yellow) complex. Key hydrogen-bond and salt-bridge residues are labeled (TARP-pep, red; ARC N-lobe, black). **c**, Left: representative PLA images; magenta puncta indicate TARP-ARC interaction. MAP2 labels neurites (blue). Scale bar, 10 μm. Right: quantification of PLA puncta. Data are mean ± s.e.m.; one-way ANOVA (Vehicle: n = 15; TARP-pep: n = 22; Rev-pep: n = 21 neurons); Tukey’s post hoc test. Significance: **** *p* < 0.0001; ns, not significant.

To bridge this gap, we designed a 21-amino-acid cell-penetrating TAT-fused peptide (TARP-pep) mimicking the TARP C-terminal motif that binds the ARC N-lobe. This design was based on the rationale that competition with the endogenous AMPAR-TARP complex for ARC binding would disrupt the TARP-ARC interaction, increase the availability of AMPAR-TARP complexes for binding to PSD-95, and thereby promote AMPAR stabilization at synapses (Fig. 1a, bottom panel). Using a combination of single-particle tracking in neuronal culture, *in vivo* electrophysiology, and behavioral approaches, we show that interfering with this interaction enhances synaptic stabilization of AMPARs, strengthens synaptic transmission in the dentate gyrus, and induces molecular signatures consistent with an LTP-like synaptic state. Importantly, manipulating this interaction also protects against social isolation-induced impairment of fear memory in mice. Together, these findings identify the TARP-ARC interface as a regulator of AMPAR diffusional trapping and suggest that targeting this mechanism may represent a strategy for preserving cognitive function under conditions of synaptic dysfunction.

## Materials and methods

### Primary neuronal cultures and transfection

For quantum dot-based single-particle tracking (QD-SPT), fluorescence recovery after photobleaching (FRAP), and proximity ligation assay (PLA) experiments, primary hippocampal neurons were prepared from E18 (embryonic day 18) embryos of both sexes obtained from 2-3-month-old timed-pregnant Wistar rats (Janvier Labs, France), as described previously^7,18^. For endogenous p-CaMKII detection, primary hippocampal neurons were prepared from E19 rat embryos of both sexes obtained from 2-3-month-old timed-pregnant Sprague Dawley rats sourced from Charles River Laboratories (Wilmington, MA), maintained under specific pathogen-free conditions. Further information regarding animal care, preparation of neuronal culture and equipment can be found in the supplementary information.

### Plasmid construction, AAV production, and neuronal application

GFP-Homer1c was generated by cloning Homer1c cDNA into pcDNA3 with eGFP fused to the N-terminus. SEP-GluA1, in which superecliptic pHluorin (SEP) is fused to the N-terminus of GluA1, was generated by cloning GluA1 cDNA into pRK5. Neurons were transfected with GFP-Homer1c or SEP-GluA1 at days in vitro (DIV) 13, and experiments were performed at DIV 14-21. The pAAV-hSyn-INTRON-iRFP-Homer1c-WPRE construct was synthesized by GenScript (Piscataway, NJ) by subcloning the cORF of DsRed-Homer1c into AAV-hSyn-mTagBFP2-CAAX2 (Addgene #236245) and replacing mTagBFP2 with iRFP670 (Addgene #45457). AAV was applied to DIV5 hippocampal neurons, medium was refreshed 2-4 days post-infection, and experiments were performed approximately 12 days after viral transduction.

### Peptide design and synthesis

A cell-permeant peptide, TARP-pep (sequence: YGRKKRRQRRR-GG-RIPSYRYR), was designed to target the TARP-ARC interface. The sequence incorporates the ARC-binding motif from TARP γ-2 (RIPSYRYR), a flexible glycine-glycine (GG) linker, and the HIV-1 Trans-Activator of Transcription (TAT) cell-penetrating domain (YGRKKRRQRRR). Two control peptides were used: Rev-pep (YGRKKRRQRRR-GG-RYRYSPIR), containing a reversed TARP motif, and TAT-pep (YGRKKRRQRRR-GG), containing only the TAT domain and linker. Rev-pep has the same charge and hydrophobic properties to TARP-pep, serving as a specific control for sequence-dependent effects. The TAT-pep lacks the TARP sequence and was used as a control to assess effects related to cell penetration. All peptides were synthesized with an N-terminal FITC-Ahx or acetyl group and a C-terminal amidation. FITC-labeled peptides were used *in vivo* to enable visualization of peptide distribution, whereas acetylated peptides were used *in vitro* to minimize potential interference from fluorophore labeling and preserve native peptide function. Custom peptides were synthesized by Synpeptide or GenScript (≥98% purity, HPLC).

### Duolink Proximity Ligation Assay (PLA)

The interaction between endogenous TARP γ-2 (stargazin) and ARC was assessed in situ using the Duolink PLA kit (Sigma, DUO92013) according to the manufacturer’s protocol. Fluorescent MAP2 (neurite marker) and PLA puncta (reporting TARP-ARC complexes) were imaged as described in the supplementary information. The analysis of PLA puncta along dendrites was performed in Fiji. Images were first converted to 8-bit, and three to five dendritic segments were blindly cropped per neuron in the MAP2 (568 nm) channel as regions of interest (ROIs). In the far-red PLA channel, a uniform intensity threshold was applied across all images to detect puncta, which were counted using the Cell Counter plugin. For each ROI, PLA density was calculated as the number of PLA puncta per dendrite length (normalized to 100 μm).

### Single-particle tracking with quantum dots (QD-SPT)

Hippocampal neurons (DIV14-21) expressing GFP-Homer1c were incubated for 2-3 h with vehicle, 1 μM TARP-pep, Rev-pep, or TAT-pep, then rinsed. Cells were blocked for 20 min and live-labeled with mouse anti-Glutamate Receptor 2 extracellular antibody (6C4) (Merck, MAB397; 1:200), followed by F(ab′)2 goat anti-mouse IgG (H+L), Qdot 655 (Thermo Fisher Scientific, Q11021MP; 1:2000; 5 min). Single-particle tracking was performed as described in the supplementary information. QDot-based trajectories were considered synaptic if colocalized with Homer1c dendritic clusters for at least five frames. To test the effect of KN-93 on the TARP-pep response, neurons were pre-treated for 1 h with 10 µM KN-93 (in DMSO) or an equal volume of DMSO vehicle. They were then incubated for 2 h with TARP-pep (in water) or water vehicle, followed by QD-SPT.

### Fluorescence recovery after photobleaching (FRAP)

Hippocampal neurons (DIV14-21) expressing SEP-GluA1 were imaged after 3-4 h incubation with vehicle, 1 μM TARP-pep, Rev-pep, or TAT-pep. FRAP experiments were performed as explained in the supplementary information. For each cell, a reference ROI (unbleached region on the same cell) and a background ROI (cell-free area) were recorded. Intensities in the bleached region (*F*_bleach_) were double-normalized to correct for background (*F*_bg_) and reference (*F*_ref_). The normalized fluorescence at time *t*, expressed as a percentage of the pre-bleach intensity (*F*_pre_), was calculated as: *F*_norm_(*t*) = [(*F*_bleach_(*t*) – *F*_bg_(*t*))/*F*_(#)_(*t*) – *F*_bg_(*t*))]/(*F*_bleach,pre_ – *F*_bg,pre_)/(*F*_ref,pre_ − *F*_bg,pre_] × 100. The mobile fraction, representing the proportion of molecules that are free to diffuse back into the bleached area, was calculated from the plateau of the normalized recovery curve as: M (%) = [(*F*_∞_ - *F*_0_) / (*F*_pre_ - *F*_0_)] × 100, where *F*_∞_ is the mean fluorescence intensity at the final plateau of the recovery, *F*_0_ is the fluorescence intensity immediately after photobleaching, and *F*_pre_ is the mean fluorescence intensity before photobleaching.

### Electrophysiology

Adult male Sprague-Dawley rats (2-3 months, 250-350 g) were used for *in vivo* medial perforant path-dentate gyrus (MPP-DG) field potential recordings. Briefly, animals were anesthetized (urethane 1.5 g/kg, *i*.*p*.), placed in stereotaxic frame, and burr holes were drilled for insertion of a stimulating electrode into the dorsomedial angular bundle and a recording electrode into the hilus of the dentate gyrus. The experimental procedure for *in vivo* infusion is described in previous works^19,20^, and information regarding the equipment used can be found in the supplementary information.

Following 20 min of baseline recordings, 1 µL of peptide (1 mM) was infused at a rate of 2.5 µL/h for 25 min, and recordings were continued for up to 2 h. Evoked field potentials were acquired and analyzed using SciWorks (version 10; DataWave Technologies). The signals were amplified, filtered (1 Hz to 10 kHz), and digitized (25 kHz). After the experiment, the rats were decapitated, and the left and right dentate gyri were removed, dissected on ice, snap-frozen in dry ice, and stored at −80°C for biochemical experiments. The procedure for co-IP can be found in the supplementary information. Electrophysiological experiments were performed by an experimenter blind to treatment.

### Detection of endogenous p-CaMKII

Endogenous p-CaMKII detection was performed as described in Benedetti *et al*.^21^, with minor modifications. Following fixation, neurons were immunolabelled with a rabbit monoclonal anti-Phospho-CaMKII (Thr286) (D21E4) antibody (Cell Signaling Technology, 12716; 1:800) and imaged as explained in the supplementary information. The identification of p-CaMKII hotspots was performed in Fiji^22^. The p-CaMKII channel was processed with Gaussian blur followed by background subtraction, where background intensity was measured from a region of interest (ROI) positioned at the dendritic center. A fixed threshold was then applied to segment p-CaMKII signal, and objects were automatically counted using Fiji’s *Analyze Particles* function. The number of objects was normalized to the surface area of the dendritic profile.

### Behavioral procedures

At 4 weeks of age, male C57BL/6 mice (Shanghai Lingchang Biotechnology Co., Ltd, China) were randomly assigned to two groups: group housing (GH) with four or five mice per cage and social isolation (SI) with only one mouse per cage. They were then raised for an additional 4 weeks before undergoing behavioral tests. Bilateral guide cannulae were stereotaxically implanted into the DG targeting the following coordinates (in mm from bregma): AP 1.95, ML ±1.5, DV –2.0, and stainless-steel obturators were inserted into the guide cannulae to prevent obstruction until peptide infusion. Following surgery, mice were allowed a two-week recovery period before fear conditioning experiments, done in accordance with previous works^23^.

Fear conditioning was conducted in accordance with a previous study^23^. Contextual and auditory fear memory were evaluated 1 h, 1 d, and 7 d after conditioning. At 1 d post-conditioning, mice received bilateral DG infusions of FITC-labeled TARP-pep or Rev-pep 3 h before the fear memory test (0.5 μl, 2 mM per side; 0.1 μl min^−1^ for 5 min). The same infusion procedure was performed once daily for 7 d to support testing at 7 d. Procedures were performed by briefly restraining mice, removing obturators, and inserting injection cannulae. The cannulae were left in place for 5 min following the infusion to permit diffusion of the peptide and minimize backflow before the obturators were replaced. All experiments involving mice were conducted in accordance with the US National Institutes of Health Guide for the Care and Use of Laboratory Animals, under an Institutional Animal Care and Use Committee (IACUC)-approved protocol, in an AAALAC-accredited facility at Shanghai Jiao Tong University School of Medicine.

Detailed description of all experimental procedures, equipment and statistical analysis can be found in the supplementary information.

## Results

### Structural modeling reveals a sequence-specific interaction between TARP-pep and the ARC N-lobe

To gain insight into the binding sites between TARP-pep and the ARC N-lobe (residues 207-278), we turned to AlphaFold 3 modeling^24^. The model predicted a high-confidence TARP-pep/ARC N-lobe interface (ARC UniProt ID: Q63053), supported by a high interface predicted TM-score (ipTM = 0.75) and low predicted aligned error (PAE) (Fig. 1b). In contrast, Rev-pep and TAT-pep models showed low confidence (ipTM = 0.49 and 0.43, respectively; high PAE; not shown). These scores suggest that the TARP-pep**/**ARC N-lobe physical interaction is sequence-specific. Subsequent binding interface analysis by PDBePISA (Proteins, Interfaces, Structures and Assemblies)^25^ revealed a compact non-covalent network wherein the core TARP-pep motif (I15-S17/Y18-R21) engages ARC residues near D210-D216/H245/N247 via multiple hydrogen bonds and R19/R21-E215/D216 salt bridges (Supplementary Table 1).

### TARP-pep disrupts TARP-ARC interaction and reduces AMPAR surface mobility at synapses

Next, we assessed the effect of TARP-pep on endogenous TARP-ARC binding in cultured hippocampal neurons using proximity ligation assay (PLA), a method of detecting proteins of interest in close proximity to each other in situ (typically within ~40 nm)^**26**^. By applying antibodies against TARP γ2 and ARC in neurons treated with TARP-pep, Rev-pep, or TAT-pep, we evaluated the effect of each peptide on the TARP-ARC interaction. Treatment with TARP-pep (1 µM, 3**-**4 h) significantly reduced the density of TARP**-**ARC interaction puncta compared to both vehicle and Rev-pep controls (one-way ANOVA, *p* < 0.0001; Fig. 1c), suggesting disruption of the endogenous TARP**-**ARC interaction.

To determine whether disruption of the TARP**-**ARC interaction enhances AMPAR diffusional trapping, we further investigated the effect of TARP-pep on AMPAR lateral diffusion within the plasma membrane using two complementary approaches: quantum dot single-particle tracking (QD-SPT) for single-molecule resolution and fluorescence recovery after photobleaching (FRAP) for ensemble behavior. AMPAR lateral diffusion, driven by Brownian motion, is highly sensitive to protein-protein interactions that constrain receptor surface mobility and can be readily disrupted, making it a sensitive readout of synaptic trapping^3,27^. AMPARs are heteromeric proteins composed of various combinations of GluA1, GluA2, GluA3, or GluA4 subunits, with GluA1-GluA2 di-heteromers being the most common in adult neurons^28,29^. We therefore investigated AMPAR surface diffusion by tracking endogenous GluA2-containing receptors using QD-SPT and recombinant SEP-GluA1 using FRAP. In this experiment, we used Rev-pep and TAT-pep as negative controls for TARP-pep, which was modified with N-terminal acetylation and C-terminal amidation to enhance stability, reduce immunogenicity, and better mimic its native form, thereby ensuring more reliable results.

QD-SPT of endogenous surface GluA2 at Homer1c-marked synapses showed that TARP-pep (1 µM, 2**-**3 h) significantly reduced the synaptic AMPAR diffusion coefficient relative to vehicle-, Rev-pep-, and TAT-pep-treated controls (Kruskal-Wallis test, *p* < 0.0001; Fig 2a), aligning with increased synaptic tethering. Consistently, FRAP of recombinant SEP-GluA1, a GluA1 subunit fused to the pH-sensitive super-ecliptic pHluorin that fluoresces primarily at the cell surface, revealed markedly slower recovery and a lower mobile fraction within dendritic spines of TARP-pep-treated neurons (1 µM, 3**-**4 h) compared with controls (one-way ANOVA; *p* < 0.0001; Fig. 2b). Together, the QD-SPT and FRAP results demonstrate that disruption of the TARP-ARC interaction markedly reduces AMPAR mobility and enhances trapping at both single-molecule and ensemble scales.

**Fig. 2.**
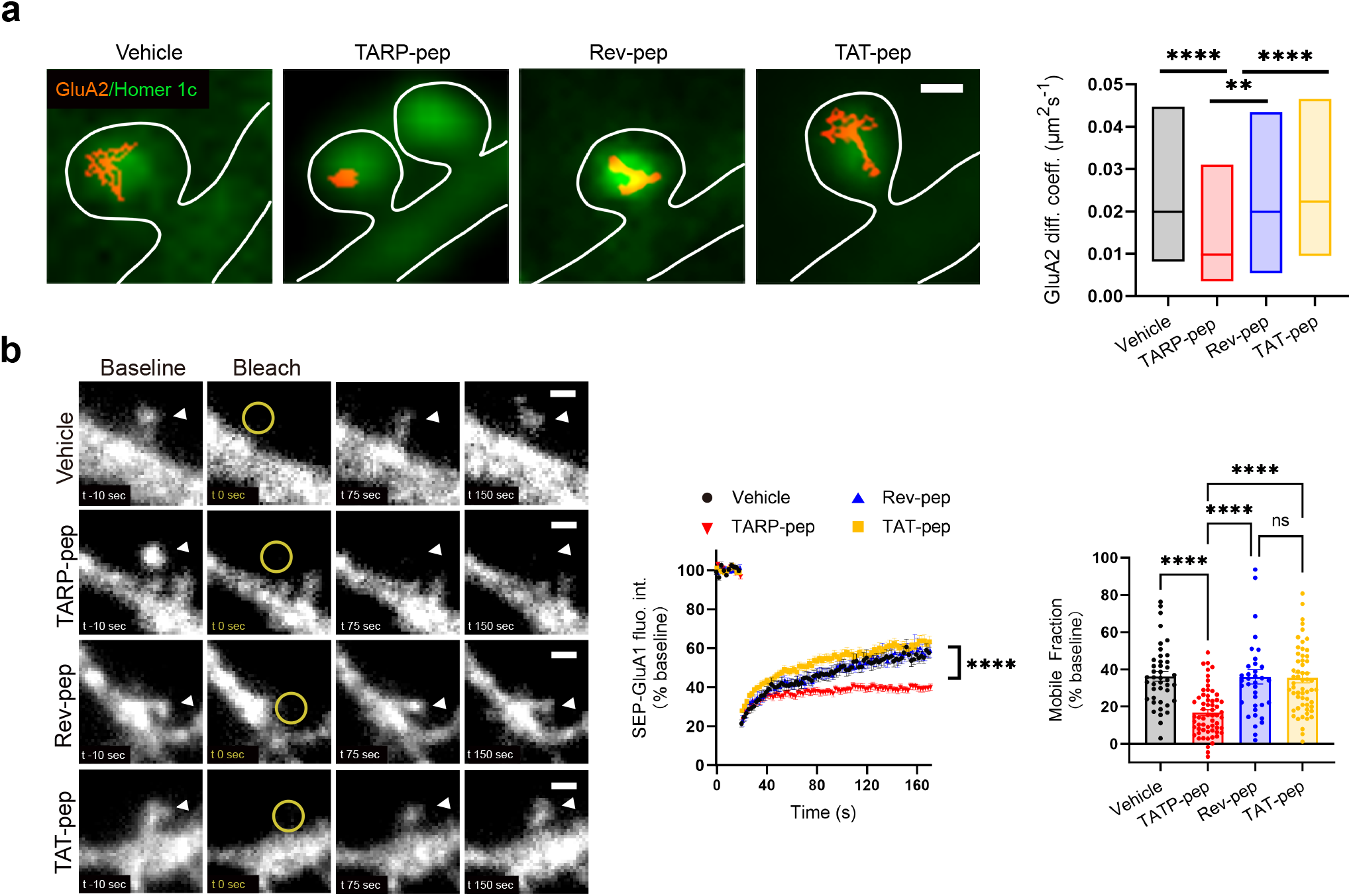
TARP-pep reduces AMPAR surface mobility at synapses across single-molecule and ensemble scales. **a**, Left: representative GluA2–QDot trajectories (red) at synapses (Homer1c puncta, green). Scale bar, 0.5 μm. Right: GluA2-AMPAR diffusion coefficients (median with 25–75% interquartile range (IOR)): Vehicle (black), TARP-pep (red), Rev-pep (blue), and TAT-pep (yellow). *n* = 344, 315, 284, 345 trajectories from 45, 51, 46, 46 synapses. Kruskal-Wallis with Dunn’s post hoc test. **b**, Left: time-lapse images before/during/after photobleaching (arrows mark synapses). Scale bar, 1 μm. Middle: FRAP recovery (normalized to pre-bleach baseline) for Vehicle (black), TARP-pep (red), Rev-pep (blue), and TAT-pep (yellow). Data are mean ± s.e.m. (*n* = 42, 51, 37, 46 synapses). Two-way RM-ANOVA showing a significant time × treatment interaction. Right: Mobile fraction (same dataset as middle), calculated as the mean of the last 20 s normalized to baseline and first post-bleach point. Data are mean ± s.e.m.; one-way ANOVA; Tukey’s post hoc test. Significance: ** *p* < 0.01, **** *p* < 0.0001; ns, not significant.

### TARP-pep enhances synaptic efficacy *in vivo* and increases hippocampal TARP–PSD-95 interaction

Having learned that competitive inhibition of the TARP-ARC interaction leads to increased synaptic trapping of AMPARs, we sought to evaluate whether these cellular effects translate to intact brain circuits. We therefore performed acute infusion of TAT-pep into the left dentate gyrus (LDG) of adult rats and measured effects on medial perforant path (MPP) evoked synaptic efficacy and changes in TARP–PSD-95 association, a key anchoring interaction that stabilizes AMPARs at postsynaptic sites^14–16^. Following baseline recording of mPP-evoked field excitatory postsynaptic potential (fEPSPs), TARP-pep or Rev-pep (1 nmol) was unilaterally infused for 25 min, followed by 2 h post-infusion recording (Fig. 3a). TARP-pep produced a gradual and robust increase in fEPSP slope relative to baseline, whereas no changes occurred in the Rev-pep control group (Fig. 3b, top and bottom left panels). At 2 hour post-infusion, fEPSPs were significantly enhanced in TARP-pep compared with Rev-pep (34.4 ± 7.7% vs. 3.8 ± 5.5%; Welch’s *t*-test, *p* < 0.01; Fig. 3b, bottom right panel). Immediately after recording, we dissected the infused LDG and the untreated contralateral right dentate gyrus (RDG), and quantified TARP–PSD-95 interaction *ex vivo* using co-immunoprecipitation (co-IP). TARP-pep, but not Rev-pep, substantially increased the TARP**–**PSD-95 interaction in the infused LDG compared to the contralateral RDG tissue (paired two-tailed *t*-test; *p* < 0.01; Fig. 3c). Together, these results demonstrate that *in vivo* infusion of TARP-pep strengthens synaptic transmission and enhances the TARP**–**PSD-95 association.

**Fig. 3.**
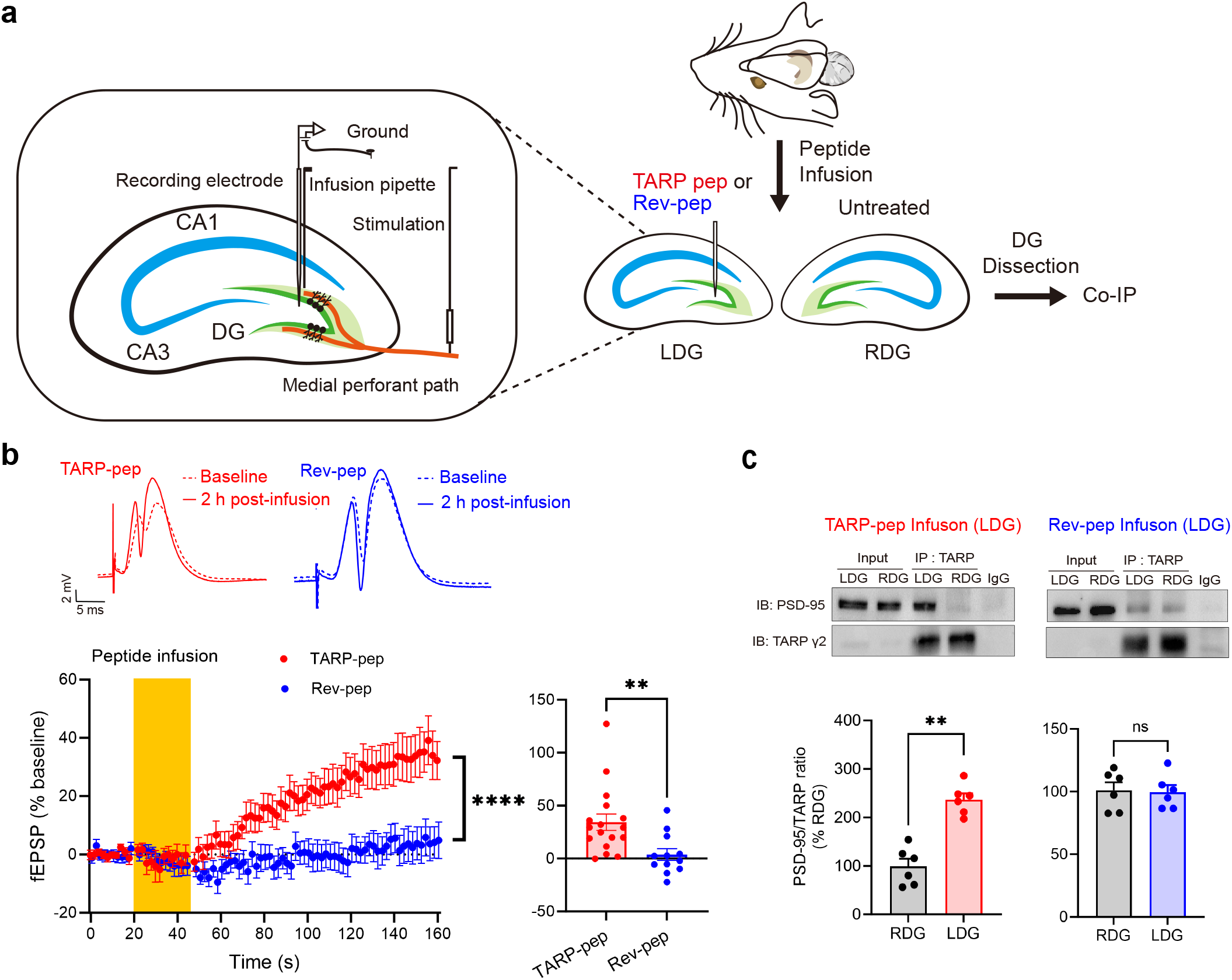
TARP-pep potentiates DG fEPSPs *in vivo* and increases TARPP–PSD-95 interaction. **a**, *In vivo* recording configuration: Stimulation of the medial perforant path, peptide infusion into stratum lacunosum-moleculare of CA1, fEPSP recording in DG hilus. DG were collected for co-IP 2 h post-infusion. **b**, Top: representative fEPSP traces at baseline (dotted) and 2 h after infusion (solid) of TARP-pep (red) or Rev-pep (blue). Scale: 5 ms, 2 mV. Bottom left: time course of fEPSP slope in LDG for TARP-pep (red, *n* = 17), and Rev-pep (blue, *n* = 12 rats); mean ± s.e.m. Two-way RM-ANOVA showing a significant time × treatment interaction. Bottom right: Mean fEPSP slope during the final 20 min (same dataset as left); mean ± s.e.m.; Welch’s *t*-test. **c**, Top: representative co-IP blots showing TARPP–PSD-95 interactions in LDG after TARP-pep (left) or Rev-pep (right) infusion versus contralateral RDG. Bottom: quantification of the TARPP–PSD-95 co-IP band intensity ratio (% of RDG). Mean ± s.e.m.; *n* = 6 mice; paired two-tailed *t*-test comparing LDG (treated) and RDG (untreated). Significance: ** *p* < 0.01, **** *p* < 0.0001.

### TARP-pep induces AMPAR synaptic trapping independent of CaMKII

To test whether TARP-pep recruits canonical long-term potentiation (LTP) mechanisms, we examined its effect on dendritic phospho-CaMKII (p-CaMKII). Neurons were pretreated with 1 µM TARP-pep, Rev-pep, or vehicle for 2 h before chemical LTP (cLTP) induction using a 30 s application of 100 µM glutamate and 10 µM glycine. TARP-pep increased baseline p-CaMKII hotspots but occluded further cLTP-induced increases, indicating that it mimics a core biochemical state of LTP (two-way ANOVA; *p* < 0.0001; Fig. 4). To test whether TARP-pep-induced AMPAR synaptic trapping depends on p-CaMKII, we applied the CaMKII inhibitor KN-93. KN-93 failed to block TARP-pep’s effect on AMPAR surface mobility at synapses, indicating that the rise in p-CaMKII is a downstream consequence of AMPAR synaptic stabilization rather than its upstream driver (Kruskal-Wallis test; *p* < 0.0001; Supplementary Fig. 1).

**Fig. 4.**
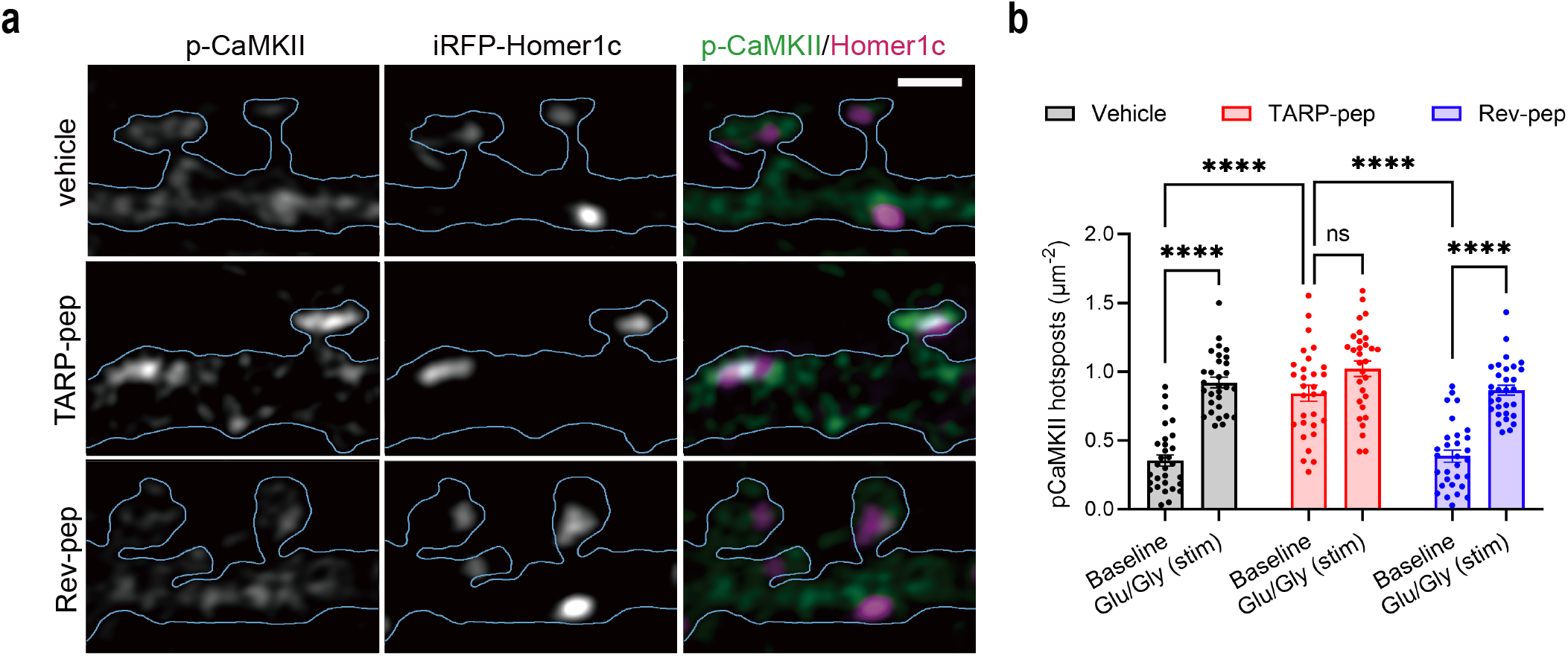
TARP-pep elevates baseline dendritic p-CaMKII and occludes cLTP-induced increases. **a**, Representative p-CaMKII immunostaining (green) following peptide treatment under baseline conditions. Magenta: Homer1c–iRFP. Scale bar, 1 μm. **b**, quantification of p-CaMKII hotspot density (number of hotspots per μm^2^ dendritic area) for Vehicle (black), TARP-pep (red), and Rev-pep (blue) under baseline conditions and following glutamate/glycine stimulation. Data are mean ± s.e.m.; two-way ANOVA (*n* = 30, 30, 31 neurons per group); Tukey’s post hoc test. Significance: **** *p* < 0.0001; ns, not significant.

### Infusion of TARP-pep prevents social isolation-induced impairment of fear memory in mice

Finally, we evaluated the *in vivo* effects of TARP-pep using a social isolation stress model, which is known to impair fear memory and reduce total levels of AMPAR subunits GluA1 and GluA2^23^. Although synaptic AMPAR levels were not directly measured, this reduction is likely to result in decreased synaptic AMPAR content. Given the critical role of AMPARs in learning and memory, we hypothesized that TARP-pep, by enhancing AMPAR synaptic trapping, could counteract social isolation-induced forgetting of fear memory.

Four-week-old male C57BL/6 mice were group-housed (GH; 4-5 per cage) or socially isolated (SI; 1 per cage) for 4 weeks. TARP-pep or Rev-pep (1 nmol) was administered bilaterally into the DG once daily for 7 days, beginning 1 day after conditioning and 3 h before each fear memory test (Fig. 5a). In these experiments, peptides were modified with an N-terminal FITC-Ahx group to enable visualization of peptide distribution and with a C-terminal amidation to enhance stability. Infusion sites were evaluated using immunostaining, confirming that the FITC-labeled peptide was in fact infused as desired (Fig. 5b). Freezing behavior was evaluated at 1 h, 1 d, and 7 d.

**Fig. 5.**
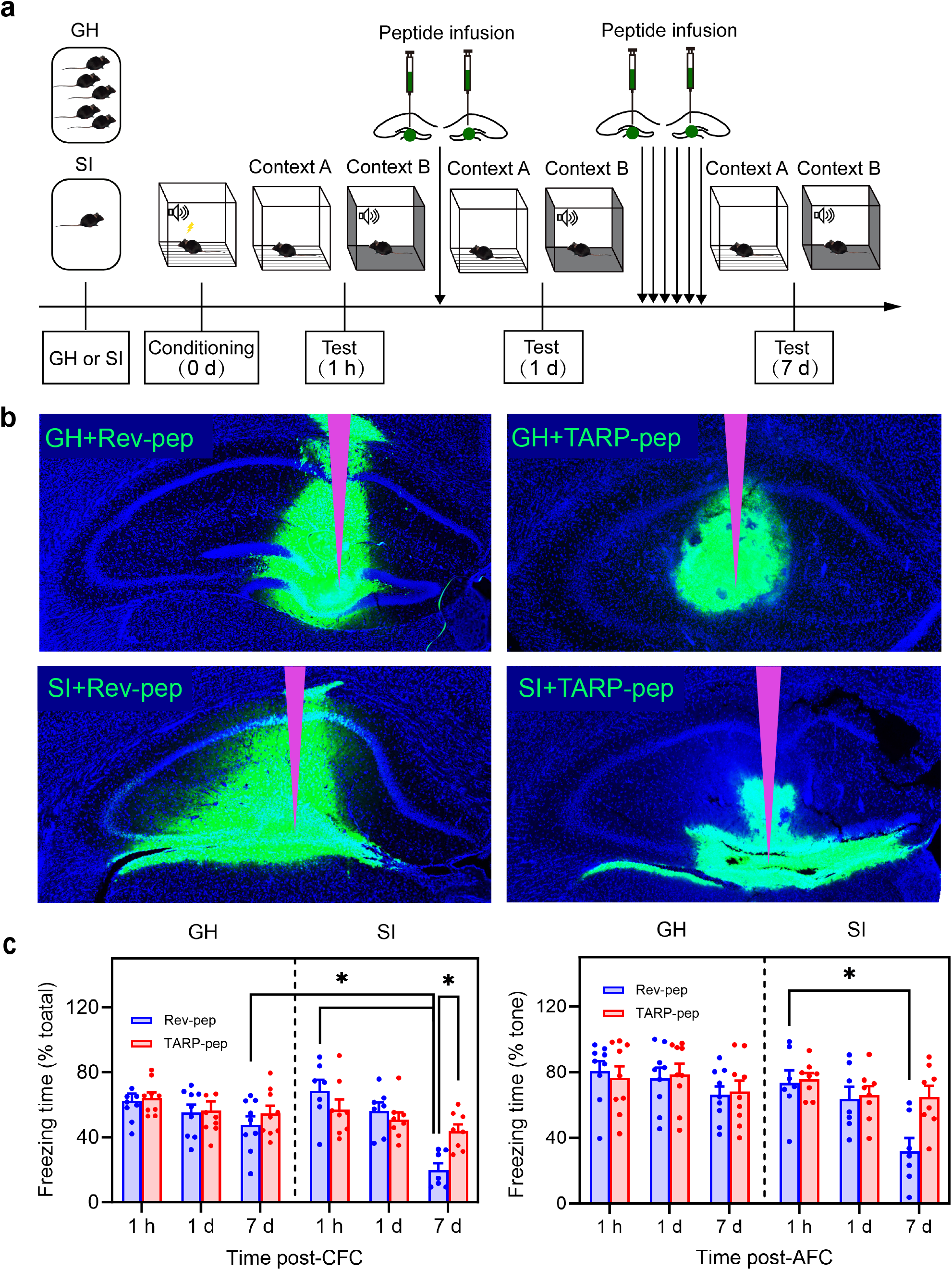
TARP-pep rescues SI-induced fear memory deficits. **a**, Experimental timeline. Four-week-old male C57BL/6 mice were group-housed (GH; 4-5/cage) or socially isolated (SI; 1/cage) for 4 weeks before fear conditioning. Memory was tested at 1 h, 1 d, and 7 d post-conditioning. TARP-pep was infused bilaterally into the DG daily for 7 d starting 3 h before the memory test on day 1. **b**, Representative immunostaining from the peptide-infusion side. Cannula placement is indicated in purple, and the peptide-infusion area is visualized by FITC-labeled peptide (green). **c**, Contextual (left, context A) and auditory (right, context B + tone) fear memory, quantified as freezing time (% of total exploration time or tone exposure), at 1 h, 1 d, and 7 d after contextual (CFC) or auditory (AFC) fear conditioning. TARP-pep (red); Rev-pep (blue). Data represent mean ± s.e.m.; *n* = 9 (GH+Rev), 9 (GH+TARP), 7 (SI+Rev), 8 (SI+TARP). Three-way RM-ANOVA with Tukey’s post hoc; Significance: * *p* < 0.05.

At one hour and one day after conditioning, there were no significant differences in contextual and auditory fear memory between GH and SI mice treated with either TARP-pep or Rev-pep. However, by day 7, SI+Rev-pep reproduced the contextual (hippocampus-dependent) and auditory (predominantly amygdala-dependent) memory deficits observed in SI mice without intervention^23^, validating Rev-pep as a negative control. In contrast, SI+TARP-pep mice showed significantly higher contextual freezing than SI+Rev-pep controls (Three-way RM-ANOVA; *p* < 0.05; Fig. 5c, left panel), whereas TARP-pep had no effect in GH mice. A similar trend was observed for auditory freezing, although this did not reach significance (Fig. 5c, right panel).

Together, these findings indicate that TARP-pep mitigates social isolation-induced impairment of hippocampus-dependent memory, consistent with an *in vivo* effect on synaptic stabilization.

## Discussion

In this study, we identify the interaction between TARPs and ARC as a critical regulatory node controlling AMPAR diffusional trapping at synapses. Using a rationally designed TAT-fused peptide that mimics the TARP C-terminal motif, we demonstrate that disruption of the TARP-ARC interaction enhances synaptic stabilization of AMPARs in cultured hippocampal neurons and strengthens perforant path-evoked synaptic transmission in the DG *in vivo*. This manipulation enhanced the interaction between TARPs and PSD-95, consistent with increased anchoring of AMPAR complexes at synapses. It also produced molecular and functional signatures of an LTP-like synaptic state, including elevated basal CaMKII phosphorylation and occlusion of further cLTP-induced CaMKII activation. Importantly, daily bilateral infusion of the peptide into the DG prevented SI-induced impairment of fear memory, linking modulation of AMPAR diffusional trapping to behavioral resilience. Together, these findings establish the TARP-ARC interface as a druggable mechanism regulating AMPAR stabilization and highlight modulation of diffusional trapping as a potential strategy for supporting synaptic and cognitive resilience.

Our findings address a long-standing challenge in neuroscience by directly modulating AMPAR synaptic trapping *in vivo* and demonstrating its therapeutic relevance in a disease-relevant model. Mechanistically, our study supports a model in which ARC-dependent regulation of the AMPAR-TARP complex is a key determinant of synaptic trapping. By competitively displacing TARP from ARC, TARP-pep shifts AMPAR-TARP complexes from a dynamic, ARC-regulated pool into a PSD-anchored state, enhancing trapping. These results provide direct *in vivo* evidence that the TARP-ARC interaction is a core physiological control point for AMPAR synaptic efficacy.

Beyond its physiological role, the pathophysiological and therapeutic relevance of the TARP-ARC interface is further revealed by the cell-permeant TARP-pep, which serves both as a mechanistic probe and as a modifiable therapeutic lead. By disrupting the inhibitory TARP-ARC interaction, TARP-pep stabilizes synaptic AMPARs, enhances synaptic transmission, and rescues SI-induced memory deficits. These results provide strong preclinical proof-of-concept that targeted enhancement of AMPAR synaptic trapping is a viable strategy for mitigating memory deficits associated with synaptic dysfunction.

The broader relevance of this strategy is supported by evidence that AMPAR synaptic trapping is impaired across models of stress/depression, Huntington’s disease, and Alzheimer’s disease, characterized by increased receptor surface mobility at synapses and weakened TARP–PSD-95 coupling^4,6,7,18,31^, as shown by us and others. Supporting this view, a study developed a “synaptic organizer” peptide that bridges presynaptic neurexins to postsynaptic AMPARs, likely stabilizing synaptic AMPARs by facilitating their binding to extracellular scaffolding proteins^32^. This peptide not only restored synaptic and cognitive function in Alzheimer’s models and but also improved motor coordination in cerebellar ataxia and spinal cord injury models^32^. Together, these findings position synaptic anchoring as a therapeutic principle across brain disorders, while our direct *in vivo* targeting of the TARP-ARC interface defines a new mechanistic route to achieve this.

TARP-Pep selectively rescues SI-induced memory deficits, with no effect in GH controls. This context-dependent effect may reflect the micromolar binding affinity of the TATpep/ARC interaction (Kd ≈ 5.3 µM)^33^, which is sufficient to relieve ARC-mediated constraint without excessively disrupting physiological signaling. Given that ARC is dysregulated in Alzheimer’s disease and stress/depression, and more broadly across multiple disease models^34–38^, this mechanism enables TARP-Pep to scale its efficacy with ARC-dependent pathology and thereby define a potential window for therapeutic selectivity. In contrast to approaches that directly modulate AMPAR, such as AMPAkines, which enhance synaptic transmission by altering receptor gating kinetics^39^, modulation of diffusional trapping at the level ARC/TARP may allow activity-dependent fine-tuning of synaptic strength, while minimizing off-target effects on AMPARs in the brain. Together, this specificity may enhance therapeutic potential while minimizing unintended effects, opening a targeted path toward interventions for synaptic vulnerability.

In conclusion, our findings introduce a novel tool to modulate AMPAR synaptic trapping and underscore the potential of targeting this mechanism as a therapeutic strategy for neurodegenerative and neuropsychiatric disorders. This work not only deepens our understanding of synaptic regulation but also paves the way for developing more targeted and effective treatments with fewer off-target effects. Although our data strongly support enhanced TARP**–**PSD-95 coupling as the main pathway, disruption of the TARP-ARC interaction may also influence other ARC-dependent processes. Secondary effects on ARC-mediated endocytosis cannot be entirely excluded and warrant further investigation using complementary assays to disentangle the relative contributions of synaptic trapping and other trafficking mechanisms. Future studies should broaden evaluation across relevant disease states, improve delivery and stability to support translation, and advance nanobodies or small molecules that precisely target the TARP-ARC interface.

## Consent for publication

All authors have read and approved the final manuscript and consent to its publication.

## Authors’ contributions

J.G., X.W., J.S., L.B., A.M., H.N., T.M., and H.Z. performed experiments and data collection: J.G. (in vivo electrophysiology, tissue collection), X.W. (behavioral tests, immunostaining), J.S. (co-IP), L.B. (assessment of CaMKII phosphorylation), A.M. (PLA), H.N. and T.M. (co-IP optimization), H.Z. (QD-SPT, FRAP). Formal data analysis was conducted by J.G. (electrophysiology), X.W. (behavior), J.S. (co-IP, FRAP, PLA), L.B. (CaMKII phosphorylation), A.P. (3D structure prediction using AlphaFold and PISA), H.Z. (QD-SPT), and W.L. & H.L. (general data analysis and interpretation). Data were validated by J.L-S. (CaMKII phosphorylation), I.G. (3D structure prediction), S.S. & N.-J.X. (behavioral experiments, immunostaining), H.Z. and C.B. (co-IP, electrophysiology, and PLA). I.G., J.L-S, S.S., N.-J.X., C.B, and H.Z. provided supervision. J.S. prepared visualizations and figures. H.Z. conceived and oversaw the entire project, including the design of the study and the peptides (TARP-pep, Rev-pep, TAT-pep). J.G. and H.Z. wrote the manuscript. All authors reviewed the manuscript and provided feedback.

## Acknowledgements

We thank Jean-Baptiste Sibarita for providing the single-particle analysis software PALM-Tracer, Endy Spriet and Hege Dale for microscopy support, and Tambudzai Kanhema Jakobsen for assistance with biochemistry. We are also grateful to François Philippe Pauzin for his input on electrophysiology and for assisting in data analysis. This work was supported by the Western Norway Regional Health Authority (Helse Vest).

Supplementary information is available at MP’s website.

---

**Supplementary Fig. 1.**
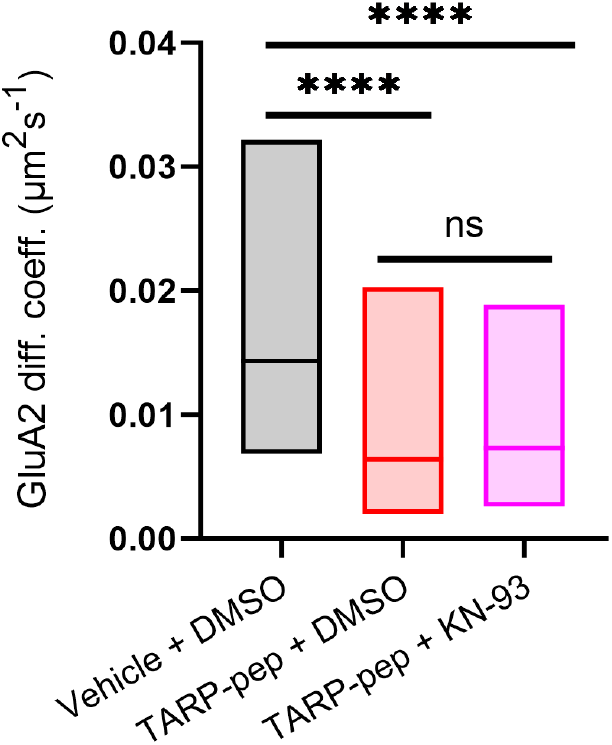
KN-93 does not prevent the TARP-pep-induced reduction in GluA2-AMPAR synaptic mobility. GluA2-AMPAR diffusion coefficients (median, 25–75% IQR) for peptide vehicle (water) + KN-93 vehicle (DMSO) (black), TARP-pep + DMSO (red), and TARP-pep + KN-93 (pink). *n* = 222, 149, and 201 trajectories. Kruskal-Wallis with Dunn’s post hoc; \*\*\*\**p* < 0.0001; ns, not significant.

**Supplementary Table 1.** Stabilizing contacts at the TARP-pep/Arc N-lobe interface. PISA-predicted contacts with bond distances (Å). The interface consists of salt bridges between TARP-pep R19/R21 and Arc N-lobe E215/D216, supported by hydrogen bonds across the TARP-pep I15–R21 segment. These contacts correspond to the complex shown in Fig. 1b.

| Hydrogen bonds |  |  |  | Salt bridges |  |  |  |
| --- | --- | --- | --- | --- | --- | --- | --- |
| # | TARP-pep | Distance (Å) | Arc N-lobe | # | TARP-pep | Distance (Å) | Arc N-lobe |
| 1 | ILE (I) 15 [N] | 2.43 | ASP (D) 210 [O] | 1 | ARG (R) 19 [NE] | 3.92 | GLU (E) 215 [OE2] |
| 2 | SER (S) 17 [N] | 2.74 | GLN (Q) 212 [O] | 2 | ARG (R) 19 [NH2] | 2.70 | GLU (E) 215 [OE2] |
| 3 | TYR (Y) 18 [N] | 2.72 | HIS (H) 245 [O] | 3 | ARG (R) 21 [NE] | 3.05 | GLU (E) 215 [OE1] |
| 4 | ARG (R) 19 [N] | 3.05 | PHE (F) 214 [O] | 4 | ARG (R) 21 [NH2] | 3.40 | ASP (D) 216 [OD1] |
| 5 | ARG (R) 19 [NH2] | 2.70 | GLU (E) 215 [OE2] | 5 | ARG (R) 21 [NH2] | 3.39 | GLU (E) 215 [OE1] |
| 6 | ARG (R) 21 [NE] | 3.05 | GLU (E) 215 [OE1] |  |  |  |  |
| 7 | ARG (R) 21 [NH2] | 3.40 | ASP (D) 216 [OD1] |  |  |  |  |
| 8 | GLY (G) 13 [O] | 2.33 | ASP (D) 210 [N] |  |  |  |  |
| 9 | ILE (I) 15 [O] | 2.87 | GLN (Q) 212 [N] |  |  |  |  |
| 10 | PRO (P) 16 [O] | 2.72 | HIS (H) 245 [ND1] |  |  |  |  |
| 11 | SER (S) 17 [O] | 2.60 | PHE (F) 214 [N] |  |  |  |  |
| 12 | TYR (Y) 18 [O] | 3.18 | ASN (N) 247 [ND2] |  |  |  |  |
| 13 | TYR (Y) 18 [O] | 3.05 | ASN (N) 247 [N] |  |  |  |  |

